# Attenuated ectopic action potential firing in parvalbumin expressing interneurons in a mouse model of Dravet Syndrome

**DOI:** 10.1101/2024.09.30.615811

**Authors:** Sophie F. Hill, Eric R. Wengert, Ethan M. Goldberg, Brian Theyel

## Abstract

Dravet syndrome is caused by heterozygous loss-of-function variants in *SCN1A*, which encodes the voltage-gated sodium channel Nav1.1. Our recent work suggests that a primary pathogenic mechanism of Dravet syndrome is impaired action potential propagation along axons of cortical parvalbumin-positive fast-spiking GABAergic interneurons (PVINs). Ectopic action potentials (EAPs) are action potentials that initiate distal to the axon initial segment. We recently demonstrated that a large proportion of PVINs fire EAPs during periods of increased excitation. Although their function remains unknown, EAPs may play a role in amplifying (when occurring in excitatory cells) and/or preventing seizures (when occurring in interneurons). Regardless of function, their generation in distal axons suggests that EAP frequency could be a useful proxy for distal axonal excitability. We hypothesized that EAPs are attenuated in PVINs from Dravet syndrome (*Scn1a*^*+/-*^) mice due to dysfunction of the distal axon. We induced EAPs in PVINs in acute brain slices prepared from male and female wildtype (WT) and *Scn1a*^*+/-*^ mice at P18-21 and P35-56, when we have previously identified axonal conduction deficits in *Scn1a*^*+/-*^ PVINs. We elicited EAPs in 17/22 (77%) of WT PVINs, including 6 (22%) that exhibited barrages of EAPs. In contrast, *Scn1a*^*+/-*^ PVINs never fired barrages (0%), and only 8/23 (34%) exhibited even single EAPs. This finding adds to the body of evidence supporting impaired action potential propagation in Dravet syndrome PVINs, and is the first evidence of impaired EAP firing in a disease model, suggesting that dysregulation of EAPs could be involved in the pathophysiology of human disease.

## Introduction

Dravet Syndrome (DS) is a severe neurodevelopmental disorder defined by treatment-resistant epilepsy onset during the first year of life, developmental delay, intellectual disability, and features of autism spectrum disorder (1). Individuals with DS have the highest known risk of sudden unexpected death in epilepsy (SUDEP) (2) and the mean age of survival is into the third decade (3, 4).

DS is caused by heterozygous loss of function variants in *SCN1A*, which encodes the neuronal voltage-gated sodium channel α-subunit Nav1.1 (5). Nav1.1 is predominantly expressed in parvalbumin positive interneurons (PVINs), which are dysregulated in DS (6–8). Haploinsufficiency of *Scn1a* causes impaired PVIN action potential generation early in development (postnatal day (P) 18-21), which is restored by P35-56 (8). However, deficits in PVIN synaptic transmission persist even at older time points (9). Taken together, these results indicate that the distal axon of PVINs is a primary locus of pathology in DS.

We hypothesized that other measures of distal axonal excitability would be affected by loss of *Scn1a*. We previously reported that PVINs exhibit high rates of action potential initiation in the distal axon, known as ectopic action potentials (EAPs, also called retroaxonal or antidromic action potentials) (10). A substantial proportion of PVINs in layer 2/3 in both primary somatosensory and orbitofrontal cortices fire EAPs, which arise after sustained action potential firing from the AIS (AIS-APs) in response to current injection. Most of the cells that fire EAPs are capable of engaging high-frequency trains that persist for seconds to minutes after stimulation, which are called “retroaxonal barrages,” “persistent firing,” “persistent barrage firing,” or simply “barrages” (11–6). Historically, EAPs have been detected in multiple cell types both in epileptic foci and in cells projecting to epileptic foci (17–20). Yet, it remains unclear whether EAPs are mechanistically involved in the pathogenesis of epilepsy.

The persistent distal axonal dysfunction in DS suggests that loss of Nav1.1 reduces axonal excitability even after somatic AP generation has normalized. The probability of EAP generation could be a proxy for the excitability of the distal axon. We hypothesize that reduced excitability of the distal axon in *Scn1a*^*+/-*^ PVINs prevents EAP firing. Here, we attempt to elicit EAPs in WT and *Scn1a*^*+/-*^ PVINs at P18-21 and at P35-56. We demonstrate that *Scn1a*^*+/-*^ PVINs are markedly less likely to generate EAPs than WT PVINs, and never exhibit barrage firing. Our findings support the idea that reduced PVIN axonal excitability is involved in DS. These are the first data demonstrating alterations in EAPs in a disease model and suggest that reduced EAP generation in inhibitory interneurons may be related to the pathogenesis of epilepsy.

## Results

To test whether loss of the voltage-gated sodium channel gene *Scn1a* impairs the generation of ectopic action potentials (EAPs), we recorded fast-spiking parvalbumin interneurons (PVINs) from acute brain slices prepared from male and female wildtype (WT) and *Scn1a*^*+/-*^ mice at P18-21 and at P35-56. We attempted to elicit EAPs by injecting increasingly large current steps lasting 600 ms until the cell entered depolarization block, a protocol that has previously been shown to reliably generate EAPs in PVINs (10). As we have previously reported, somatic evoked spiking (AIS-APs) was impaired in PVINs from P18-21 *Scn1a*^*+/-*^ compared to WT (**Fig S1**) (8, 9). At P35-56, we observed no change in the excitability of *Scn1a*^*+/-*^ PVINs compared to WT, consistent with our previous observations that excitability is restored at this time point (**Fig S1B**) (8, 9).

### Reduced EAP frequency in *Scn1a*^*+/-*^ PVINs

At both P18-21 and P35-56, 7/10 WT PVINs exhibited ectopic spiking (**Fig 1A-B**). PVINs from *Scn1a*^*+/-*^ mice were less likely to fire EAPs at both time points (P18-21: *p* = 0.1789; P35-56: *p* = 0.0414; all ages: *p* = 0.0067; *p*-values from Fisher’s exact tests) (**Fig 1A-B**).

**Fig. 1.**
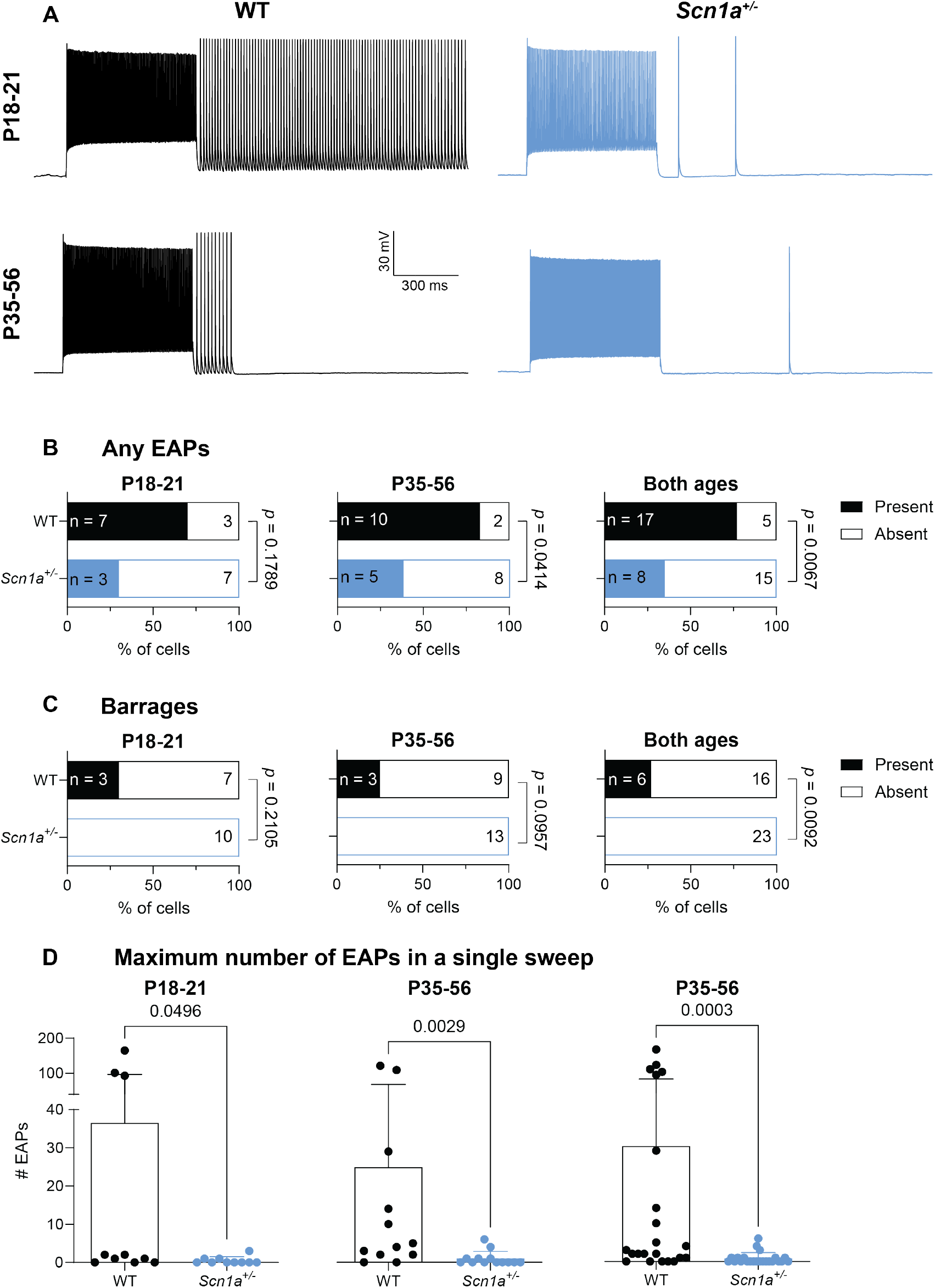
Reduced frequency of ectopic action potentials in *Scn1a*+/-PVINs. **A**, Representative example sweeps from the EAP initiation protocol. **B & C**, Proportion of WT and *Scn1a*^*+/-*^ PVINs that exhibited any ectopic action potentials (B) or barrage firing (C). P-values indicate significance of Fisher’s exact tests. **D**, Maximum number of EAPs in a single sweep. Each point represents a single cell. P-values indicate significance of Mann-Whitney U-tests.

Interneurons have previously been shown to fire trains of dozens to hundreds of EAPs (termed “barrages”) in response to somatic stimulation (**Fig. 1A, top left**) (10, 11). At P18-21, 30% of all WT PVINs and approximately half of the PVINs that exhibited any EAPs fired prolonged barrages. In comparison, no *Scn1a*^*+/-*^ PVINs exhibited barrage firing (*p* = 0.2105, Fisher’s exact test) (**Fig 1A&C, Table 1**). The same trend was observed at P35-56 (*p* = 0.0957, Fisher’s exact test) and was statistically significant when PVINs from both ages were pooled (*p* = 0.0092, Fisher’s exact test) (**Fig 1C, Table 1**).

**Table 1.**
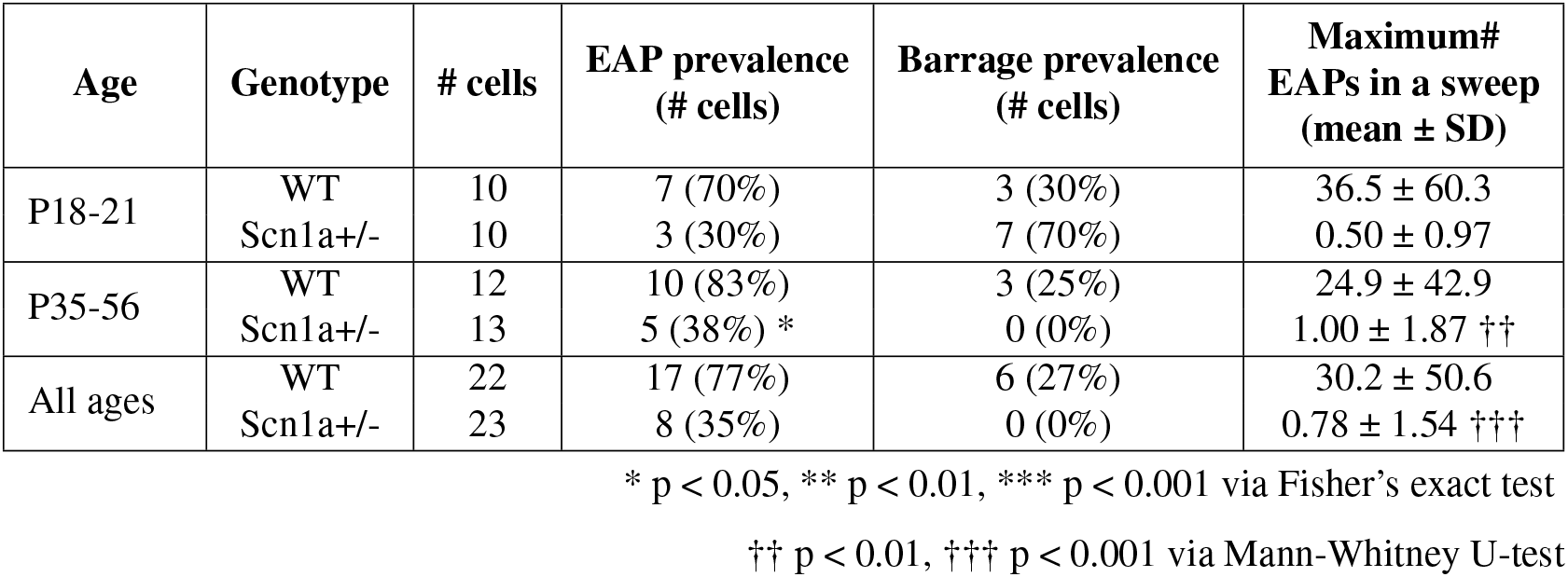
EAP prevalence.

**Table 2.**
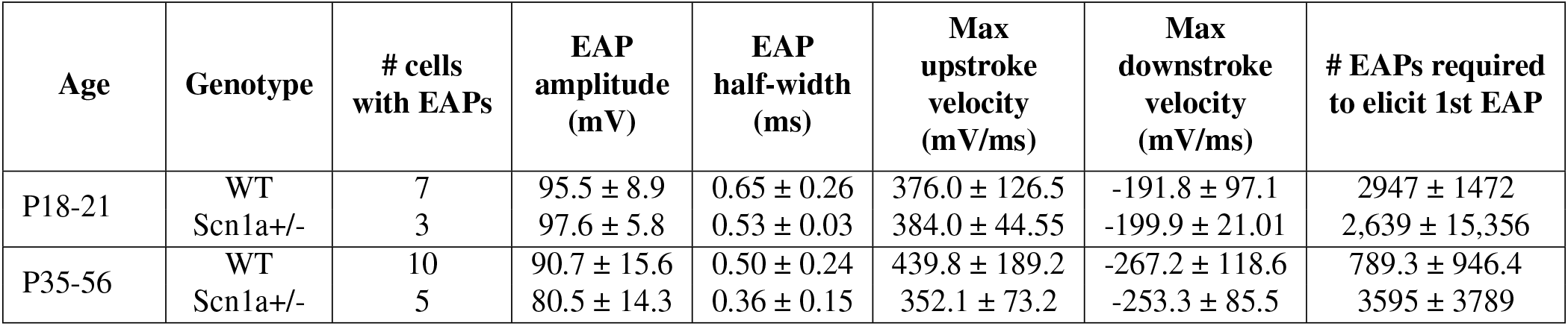
EAP properties.

For each cell, we quantified the maximum number of EAPs observed in a single sweep (**Fig 1D, Table 1**). At every age, *Scn1a*^*+/-*^ PVINs exhibited fewer EAPs (P18-21: *p* = 0.0496; P35-56: *p* = 0.0029; all ages: *p* = 0.0003; *p*-values from Mann-Whitney U-tests). Of the *Scn1a*^*+/-*^ PVINs that exhibited any EAPs, most (5/8) did not fire more than 1 EAP per sweep (**Fig 1D**).

### EAP waveform in *Scn1a*^*+/-*^ PVINs

We hypothesized that heterozygous loss of Nav1.1 could affect the properties of EAPs. The mean amplitude of EAPs from *Scn1a*^+/-^ PVINs was not significantly different from WT at either P18-21 (*p* > 0.9999, Mann-Whitney U-test) or P35-56 (*p* = 0.2544, Mann-Whitney U-test) (**Fig 2A, Table 1**). Similarly, there was no change in the maximum upstroke (P18-21: *p* = 0.4833; P35-56: *p* = 0.6354, Mann-Whitney U-tests) or downstroke velocity (P18-21: *p* = 0.3833; P35-56: *p* = 0.7679, Mann-Whitney U-tests) (**Fig 2B&D, Table 1**). We also found no difference in EAP half-width at either time point (P18-21: *p* = 0.3833; P35-56: *p* = 0.5704, Mann-Whitney U-tests). These findings are consistent with previous studies of AIS-APs, in which heterozygous loss of *Scn1a* causes no or only minor changes in the AP waveform (7–9). The number of AIS-APs required to elicit the first EAP was higher in *Scn1a*^*+/-*^ PVINs, although this effect was not statistically significant (P18-21: *p* = 0.3833; P35-56: *p* = 0.7679, Mann-Whitney U-tests, **Fig 2E, Table 1**).

**Fig. 2.**
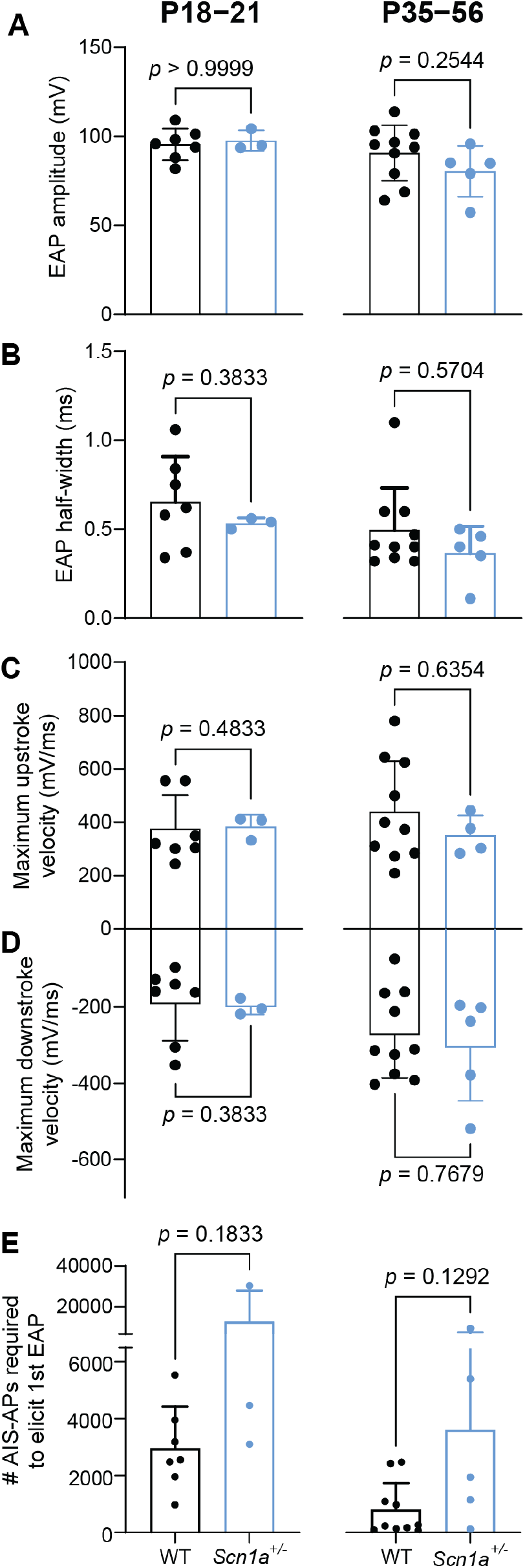
EAP properties are unchanged by loss of *Scn1a*. **A**, EAP amplitude. **B**, EAP half-width. **C**, Maximum EAP upstroke velocity. **D**, Maximum EAP downstroke velocity. **A-D** were calculated from the first EAP in each cell. **E**, Number of AIS-APs required to elicit the first EAP in each cell. P-values indicate significance of Mann-Whitney U-tests.

## Discussion

We have demonstrated a deficiency in EAP generation in PVINs with haploinsufficiency of the voltage-gated sodium channel Nav1.1. Our results are consistent with a view of Dravet syndrome as an interneuron “axonopathy”, in which the primary pathogenic mechanism is persistent loss of excitability in the distal axon of PVINs, despite restoration of AIS-AP generation by P35-56. To the best of our knowledge, this is also the first evidence linking impaired EAP generation and disease.

EAPs are generally underexplored, but recent studies suggest that a multitude of mouse neuronal types exhibit EAPs in response to repeated stimulation (10, 21). The mechanism of EAP generation is unknown, but there is some evidence that gap junctions and calcium signaling are required for barrage firing (11, 12). There is also evidence suggesting that hyperpolarization-activated cyclic nucleotide-gated (HCN) channels and astrocytes may be implicated (15, 16). EAPs have been extensively described in crustaceans(22–25), peripheral nerve of the cat (26), and models of epilepsy in mice, rats, and cats

EAP detection in epilepsy is of unclear significance: do EAPs participate in the disease state, protect against seizures, or are they merely observed as a result of increased activity? It is possible that a pathological increase in EAP generation in excitatory cells (or decrease in inhibitory cells) could shift the overall balance of excitation and inhibition, yielding seizures. Since EAPs are only observed after periods of increased activity, any shift in the propensity of inhibitory vs. excitatory cells to fire EAPs could amplify or inhibit seizure propagation. There is some evidence that EAPs can be synaptically transmitted (13). However, EAPs may also arise in order to drive homeostatic mechanisms in the soma in response to repeated stimulation.

While no direct comparisons have been made, several studies suggest that EAPs are robustly generated in mammalian inhibitory interneurons, whereas EAP generation in excitatory cells tends to be weaker (10, 11, 13, 14, 16, 21). We found, for instance, that >90% of PVINs in layer 2/3 mouse orbitofrontal cortex (OFC) and 100% of upper-layer primary somatosensory cortex PVINs are capable of generating EAPs (21). This contrasts with our finding that 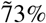 of OFC pyramidal cells are capable of firing EAPs. Perhaps more compelling was our finding that OFC PVINs fired an average of 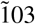 EAPs when maximally engaged, whereas pyramidal cells only fired an average of 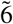. Thus, we and others have theorized that EAPs could serve as an overall ‘brake’ that is engaged during hyperexcitability. However, other studies have suggested that smaller proportions of inhibitory cells fire EAPs. For example, Imbrosci et al. (2015) detected EAPs in only 1/24 (4%) of fast-spiking (presumed PVINs) in the visual cortex (27). Suzuki et al. (2014) demonstrated that 24% of fast-spiking cells fired EAPs, whereas 85% of neurogliaform cells fired EAPs (13). In the present study, 17/22 (77%) of WT somatosensory PVINs fired EAPs. While the source of the variability in EAP generation probability remains unknown, differences in experimental conditions, interregional differences, animal strain, age, or ascertainment bias (e.g. reporting barrages of EAPs but not single EAP generation) could be explanatory (10).

Here, we demonstrate that PVINs are less likely to fire EAPs in a mouse model of Dravet syndrome. PVINs are the primary driver of the DS phenotype (7). We have previously shown impaired PVIN AIS-AP generation in the developmental window between P16-21 (8). At older time points, PVIN AIS-AP generation is partially restored, but deficits in axonal transmission persist (8, 9, 28). Our data demonstrate an additional deficiency manifested by this persistent distal axonal dysfunction: an impaired ability to generate EAPs in PVINs, which could serve as highly potent sources of neocortical inhibition. The deficiency in EAP generation is likely caused directly by reduced voltage-gated sodium channel activity in the distal axon.

The role of EAPs in epilepsy is still controversial, but our findings are consistent with a view of EAPs as neuroprotective. Reduced EAP generation could yield a significant reduction in overall inhibitory output both before and during seizures, and thus could blunt a source of negative feedback that would normally prevent development of seizures during epochs of increased activity or terminate seizures once they have started. It is unclear whether our finding is specific to *Scn1a*^*+/-*^ mice, or whether reduced EAP generation is a common feature of sodium channelopathies or all genetic epilepsies.

Epilepsy in Dravet Syndrome is typically treatment resistant. Here we have demonstrated a significant deficiency in the ability of potent neocortical inhibitory interneurons to general EAPs. This represents the first finding that abnormalities in EAP generation are linked to pathology and identifies a novel physiological target in treatment discovery.

## ACKNOWLEDGEMENTS

We acknowledge Dr. Xiaohong Zhang and Emily Hoddeson for technical support. This work was funded by NIH NINDS R01 NS110869 to E.M.G., a Dravet Syndrome Foundation Postdoctoral Fellowship to S.F.H., and a Dravet Syndrome Foundation Research Grant to B.T.

