## Supplementary materials for "Attenuated ectopic action potential firing in parvalbumin expressing interneurons in a mouse model of Dravet Syndrome"

Fig S1

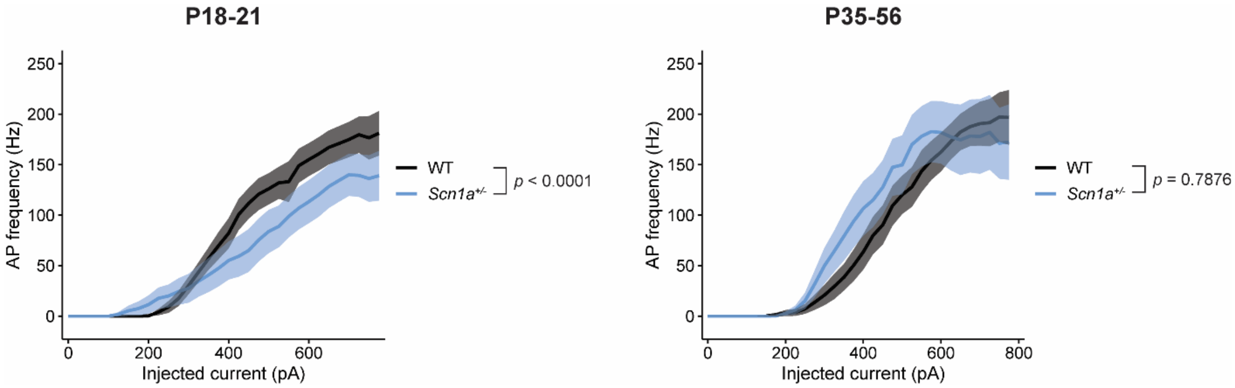


**Supplementary Figure 1. Developmental impairment of *Scn1a*$^{+/-}$ PVINs.** Input-output curves for WT and *Scn1a*$^{+/-}$ PVINs at P18-21 and P35-56. P-values indicate the significance of the main effect of genotype in two-way ANOVA tests.

Fig S2

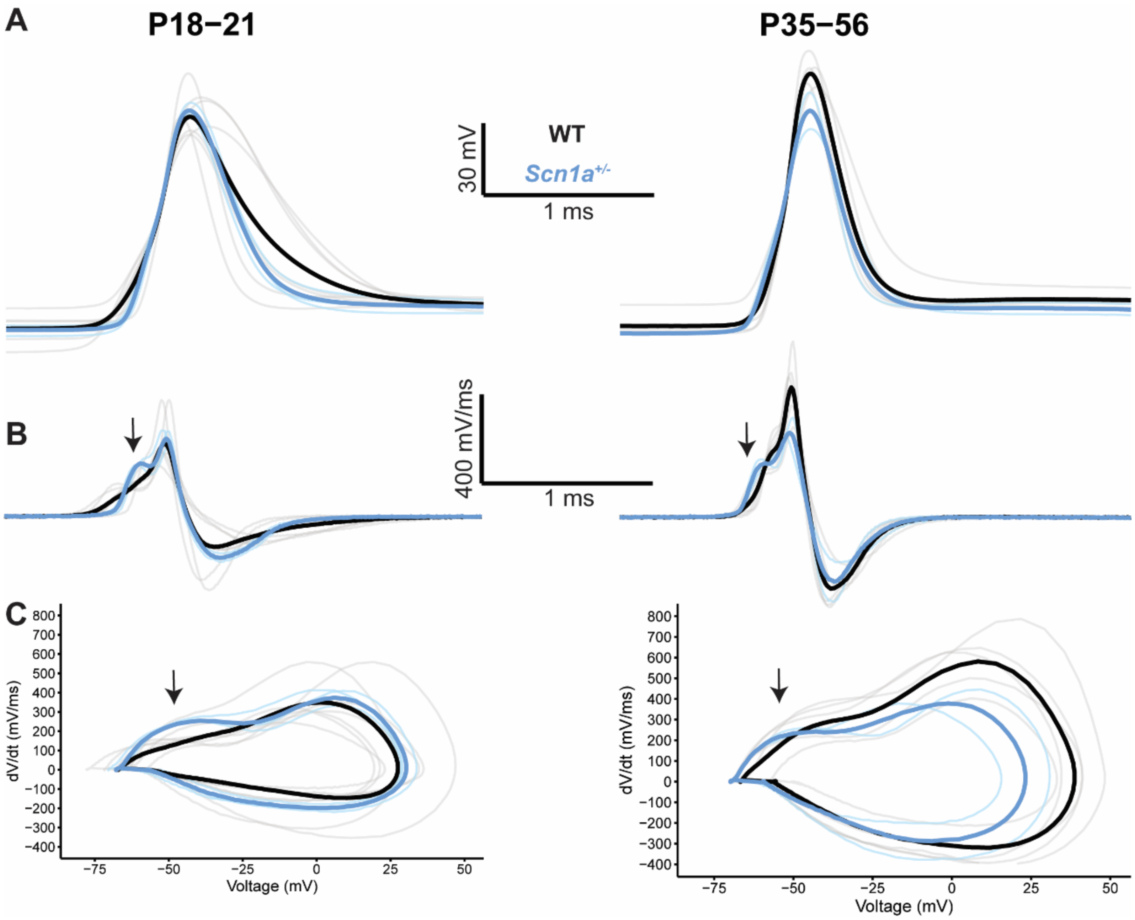


**Supplementary Figure 2. Prominent first hump in EAPs from *Scn1a*$^{+/-}$ PVINs. A,** Overlay of the first EAP in every cell and the genotype averages. **B**, Overlay of the first derivative of the voltage (mV/ms) from the first EAP in every cell, and the genotype averages. Arrows indicate a more prominent first inflection point in the *Scn1a*$^{+/-}$ EAPs. **C,** Phase plots (first derivative as a function of voltage) of EAPs overlaid with the genotype averages. Again, arrows indicate a more prominent first inflection point in the *Scn1a*$^{+/-}$ EAP phase plots. EAPs were aligned at the point where the first derivative of the voltage exceeded 10 mV/ms (EAP threshold).
